# Single-cell Transcriptional Analysis of the Cellular Immune Response in the Oral Mucosa of Mice

**DOI:** 10.1101/2023.10.18.562816

**Authors:** P Cantalupo, A Diacou, S Park, V Soman, J Chen, D Glenn, U Chandran, D Clark

## Abstract

Periodontal health is dependent on a symbiotic relationship of the host immune response with the oral microbiota. Pathologic shifts of the microbial plaque elicit an immune response that eventually leads to the recruitment and activation of osteoclasts and matrix metalloproteinases and the eventual tissue destruction that is evident in periodontal disease. Once the microbial stimulus is removed, an active process of inflammatory resolution begins. The goal of this work was to use scRNAseq to demonstrate the unique cellular immune response across three distinct conditions of periodontal health, disease, and resolution using mouse models. Periodontal disease was induced using a ligature model. Resolution was modeled by removing the ligature and allowing the mouse to recover. Immune cells (Cd45+) were isolated from the periodontium and analyzed via scRNAseq. Gene signature shifts across the three conditions were characterized and shown to be largely driven by macrophage and neutrophils during the periodontal disease and resolution conditions. Resolution of periodontal disease was characterized by the differential regulation of unique gene subsets. Clustering analysis characterized multiple cellular subpopulations within B Cells, macrophages, and neutrophils that demonstrated differential expansion and contraction across conditions of periodontal health, disease, and resolution. Interestingly, we identified a transcriptionally distinct macrophage subpopulation that expanded during the resolution condition and demonstrated an immunoregulatory gene signature. We identified a cell surface marker for this resolution-associated macrophage subgroup (Cd74) and validated the expansion of this subgroup during resolution via flow cytometry. This work presents a robust immune cell atlas for study of the immunological changes in the oral mucosa during three distinct conditions of periodontal health, disease, and resolution and it improves our understanding of the cellular and molecular markers that characterize health from disease for the development of future diagnostics and therapies.

## Introduction

Periodontal health is dependent on a symbiotic balance of the host immune response with the oral microbiota. When pathologic shifts of the microbial plaque occur, qualitative and quantitative changes of the host cellular immune response are evident^1^. Dysbiotic shifts in the immune response may precede clinical signs of disease^2^. The initial innate immune response in periodontal disease is characterized by neutrophil and macrophage infiltration followed by an adaptive immune response with evident T cell expansion^3,4^. Eventual recruitment and activation of osteoclasts and matrix metalloproteinase activity leads to the osteolytic inflammatory response and tissue destruction evident in periodontal disease^5^.

Removal of the pathologic microbial plaque eliminates the immune response stimulus, and the process of inflammatory resolution occurs to promote the return to tissue homeostasis^6^. A timely and complete resolution of inflammation is necessary to limit the tissue damage. Resolution of inflammation is an active process beyond just the cessation of the pro-inflammatory signaling. The active process of resolution consists of the production of mediators that inhibit further immune cell recruitment, upregulation of apoptosis and efferocytosis and the removal of apoptotic cells and debris, and initiation of the cascade of regenerative processes to repair the damaged tissue^7-9^.

Resolution of inflammation is also characterized by phenotypic changes of immune cells. Phenotypic changes of macrophages during periodontal health and disease have been previously characterized^10^. Macrophages phenotype has been generally described as pro-inflammatory (M1) and anti-inflammatory (M2). Advances in the field and incorporation of high throughput transcriptional analysis have led to the characterization of many diverse macrophage phenotypes that are tissue, disease, and developmental origin specific^11-13^. A better understanding is needed of the diverse macrophage phenotypes during periodontal health and disease.

The goal of this work was to demonstrate the unique cellular immune response across three distinct conditions of periodontal health, disease, and resolution in mice using single-cell RNA sequencing (scRNAseq). We present the unique transcriptional signatures and cellular subgroups that characterize each condition. Interestingly, we identified a transcriptionally distinct macrophage subgroup that expanded during the resolution conditions and demonstrated an immunoregulatory gene signature. We identified a cell surface marker for this resolution-associated macrophage subgroup and validated the expansion of this subgroup during resolution via flow cytometry. These findings begin to identify markers of inflammatory resolution in periodontal disease to determine earlier and more accurate markers of disease progression versus resolution.

## Methods

### Periodontal disease mouse model

The animal procedures were approved by the University of Pittsburgh Institutional Animal Care and Use Committee and were performed in accordance with the guidelines of the US National Institutes of Health for the care and use of laboratory animals and the ARRIVE checklist. Male C57BL/6 mice were used for all experiments at the age of 4 months. Three groups of mice were compared in each analysis: healthy controls (control), periodontal disease induction (disease), and periodontal disease resolution (resolution). Periodontal disease was induced using the ligature method as described previously^14^. Mice were first anesthetized with an intraperitoneal injection of a 1:1 solution of dexmedetomidine and ketamine. A 6.0 silk suture was tied in a subgingival position around the second maxillary molars bilaterally. The sutures remained in place for 7 days to induce a local osteolytic inflammatory response and characteristic periodontal disease phenotype. In the disease group, mice were euthanized after 7 days and tissues were isolated for analysis. In the resolution group, the ligature was removed after 7 days, and mice were allowed to recover for an additional 5 days before they were euthanized.

### Bone loss measurements

Periodontal disease severity was evaluated by quantifying the extent of linear alveolar bone loss in the maxilla of mice. Mice from control, disease, and resolution groups were compared (n=7/group). One half of the maxilla containing the 3 molar teeth was isolated, defleshed, and placed in 30% hydrogen peroxide overnight, and then fixed in 70% ethanol. Magnified images of the maxilla were captured using a dissecting microscope (Leica M165 FC, Leica Microsystems Ltd). One image of the buccal aspect and one image of the palatal aspect of the maxilla were captured for each sample. Measurement calibrations were made possible by including a ruler in each image placed adjacent to the sample. Linear bone loss was measured from the cementoenamel junction (CEJ) to the alveolar bone crest (ABC) at 6 sites per tooth (3 sites on the buccal and 3 sites on the palatal surface of each tooth). Bone loss measurements were made on the images using ImageJ. All measurement sites were averaged per animal.

### Quantitative Real-time Polymerase Chain Reaction of gingival mucosa

The extent of inflammation locally in the gingiva of the mice was evaluated by quantifying inflammatory cytokine gene expression via quantitative real-time polymerase chain reaction (qRT-PCR). Mice from control, disease, and resolution groups were compared (n=7/group). Gingiva was isolated from one half of the maxilla. The isolated gingiva was prepared for qRT-PCR analysis. Briefly, gingiva was homogenized in Trizol and RNA was isolated. cDNA was reverse transcribed using Superscript III (Invitrogen). qRT-PCR was performed with SYBR Green and the following primers;

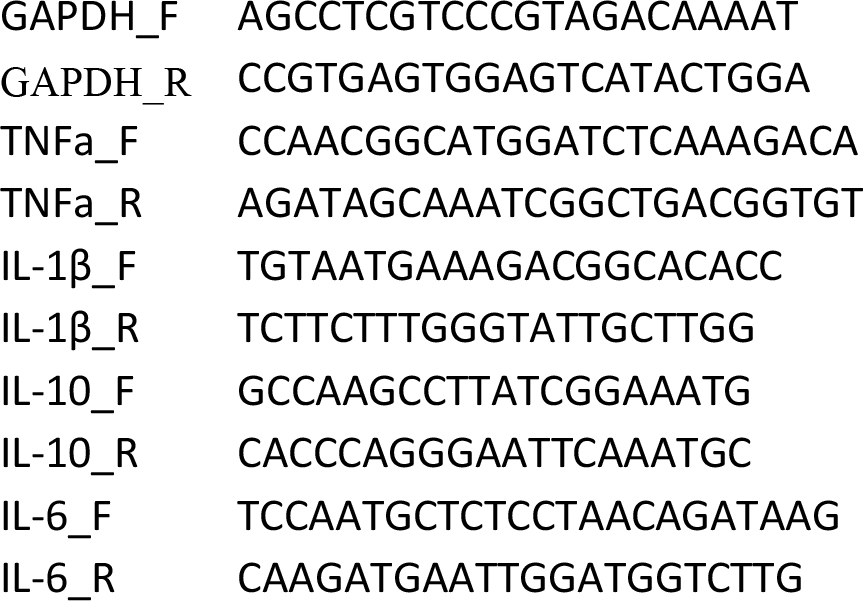

### Flow cytometry

Both halves of the maxilla were isolated from mice in the control, disease, and resolution groups (n=6/group). Maxilla were disassociated manually and then strained using a 100μm cell strainer. Tissues was further digested with collagenase type 1 (0.2mg/ml, Worthington Lakewood, NJ) for 1 hour at 37°C. Cells were washed, collected by centrifugation, and resuspended in incubation buffer (0.5% BSA in PBS). Isolated cells were blocked for 10 min in 10% rat serum. To detect dead cell, cells were stained with Fixable Viability Dye (BD Biosciences Franklin Lakes, NJ). Cell were also stained with directly conjugated fluorescent antibodies (Rat anti-mouse): CD45 (clone 30-F11), MHC Class II (M5/114.15.2), Ly6C (AL21), Ly6G (clone 1A8), F4/80 (T45-2342), CD11b (clone M1/70) (BD Biosciences, Franklin Lakes, NJ), and Cd74 (clone 829706) (R&D Systems, Minneapolis, MN). Fluorescence minus one controls were used to gate for background staining. Cells were analyzed and/or sorted on a FACSAria (BD Biosciences, San Jose, CA). FlowJo Software 9.6 (Treestar, Ashland, OR) was used for analysis.

### Statistical analysis

The linear bone loss and cell quantification were calculated per sample and presented as mean ± standard deviation (SD). Analysis of variance (ANOVA) was utilized to first test for significant differences across all groups. Further, between-group differences were analyzed using a 2-tailed t test. Quantitative real-time polymerase chain reaction (qRT-PCR) used technical triplicates and the mean QT value was calculated. mRNA expression (2^-ΔCT) was calculated and presented as mean ± standard error of the mean (SEM) and analyzed using an ANOVA followed by between-group comparisons using a 2-tailed t test. Significance for all analysis was determined at p< 0.05. All statistical analysis was performed using GraphPad Prism v.7 software (GraphPad Software, Inc.)

### Single cell RNA-sequencing

Cells were isolated from the periodontium of mice in control, disease, and resolution conditions (n=3/group). Viable CD45+ cells were isolated via flow cytometry as described above. The isolated cells were prepared in a single cell suspension and library preparation was performed using the Chromium Single Cell 3’ Reagent Version 3 Kit (10X Genomics). Each sample was loaded and run on separate lanes. Droplets were subjected to reverse transcription and then cDNA amplified. The prepared libraries were sequenced on an Illumina HiSeq4000. The mouse reference genome, mm10, was downloaded from 10X Genomics (https://cf.10xgenomics.com/supp/cell-exp/refdata-gex-mm10-2020-A.tar.gz). Raw reads were processed using the reference genome with Cell Ranger (v6.1) to generate raw UMI count tables.

### Single cell RNA-seq analysis

Seurat version 4 was used for analysis^15^. The percentage of mitochondrial reads per cell was determined using PercentageFeatureSet. Then, cells were removed according to the following metrics: < 300 features expressed, UMI counts < 1,000 or > 50,000, or > 10% mitochondrial reads. Genes were removed that were not expressed in at least 3 cells. After filtering, the number of cells remaining ranged from 892 to 2453 and the number of genes remaining was 20,416. The raw UMI counts were normalized using NormalizeData. The Seurat ‘cc.genes.updated.2019’ human gene symbols were converted to mouse symbols using the homologene R package and subsequently used to determine the cell cycle phase of each cell with CellCycleScoring. Then, variable genes (HVG) were determined using the ‘vst’ method. To determine cell cycle associated HVGs, we used getVarianceExplained from the R package scater which identified 294 genes that we removed from the HVG list (1706 HVGs remained). Then the HVGs were scaled with regression on ‘nCount_RNA’ and used to determine a PCA. We determined the number of PCA dimensions to use by identifying the PCs accounting for 75% of the cumulative PCA standard deviation (PCs 1 to 29). These PCs were used to generate a UMAP using RunUMAP and clusters were determined with FindClusters using default parameters.

#### Subclustering

The expression of immune cell type markers was visualized with FeaturePlot and used to annotate cell types to each cluster. Three clusters were removed without transcriptional signatures of known immune cell types. Seurat was used to analyze and recluster the remaining cells (ranging from 856 to 2396 cells per sample) similar to the methods above. Briefly, 326 HVGs were found to be associated with the cell cycle and removed (1674 HVGs remained). After scaling and PCA, the first 29 PCs were used to generate a UMAP and determine cell clusters.

#### Markers

Cell type markers were determined using FindAllMarkers. Differentially expressed genes between conditions per cell type or across all cells (aka. global) was determined using FindMarkers. The defaults were used for the two functions except that the ‘min.pct’ parameter was increased to 0.25. Heatmaps of the top 5 or 25 genes were generated with DoHeatmap. To generate volcano and boxplots, cell type markers and global DEG were filtered to remove ribosomal genes.

#### Gene Ontology Analysis

The resolution vs disease differentially expressed genes for Macrophages and Neutrophils were filtered to remove ribosomal genes and on adjusted P-value < 0.05 to obtain lists of significantly up and downregulated genes. These gene lists were uploaded to Enrichr (https://maayanlab.cloud/Enrichr/) to obtain the GO Biological Process 2023 regulated pathways. Barplots were generated with ggplot2 using the -log10(P-value) and colored by combined score.

#### CellChat

CellChat version 1.6.1 was used to infer cell-cell communication between cell types^16^. The CellChat vignette for a single dataset was followed to generate a CellChat object for each condition (control, disease, and resolution) using the “Secreted Signaling” subset of the mouse database. To compare two conditions, we used the multi-sample vignette. A circle plot was generated using netVisual_diffInteraction to show the differential number of interactions between cell populations. Bubble plots were created using netVisual_bubble to show the upregulated ligand-receptor pairs between two conditions.

#### Data Visualizations

Several R packages were used to generate plots. Volcano plots were built using EnhancedVolcano (v.1.16.0), heatmaps were generated using pheatmap (v1.0.12) and ggplot2 (v3.3.6) was used to generate boxplots and percent stacked barplots.

## Results

### Validation of periodontal disease induction and resolution model

We utilized a ligature model in mice to study the immune response across conditions of periodontal health (control), periodontal disease (disease), and resolution from periodontal disease (resolution) (n=7/group). To induce disease, ligatures were tied around the second maxillary molar for 7 days. To analyze resolution, ligatures were removed after the disease induction period and the animal was allowed to recover for 5 days (Fig. 1A). Flow cytometry analysis of immune cells (Cd45+) infiltrating into the periodontium showed that immune cells peaked in the disease group and returned to near baseline levels in the resolution group (Fig. 1B). Linear bone loss measures of the maxillary alveolar bone showed that bone loss was significantly greater in the disease group compared to the others. In the resolution group, there was a significant gain in bone level compared to the disease group, but bone levels did not return to baseline and there was still significantly greater bone loss compared to control group (Fig. 1C,D). We then used qPCR analysis to evaluate inflammatory gene expression within the periodontium. Expression of inflammatory cytokines Il-1β and Il-6 was significantly increased in the disease group compared to control. Reduction of the inflammatory genes was observed in the resolution group. Anti-inflammatory gene Il-10 remained significantly elevated in the resolution group compared to control (Fig. 1E). Taken together, we validated our model of periodontal disease by showing that the resolution group was associated with alveolar bone gain and decreased inflammatory cytokine expression compared to the disease group but without a complete return to baseline conditions observed in the control group.

**Figure 1.**
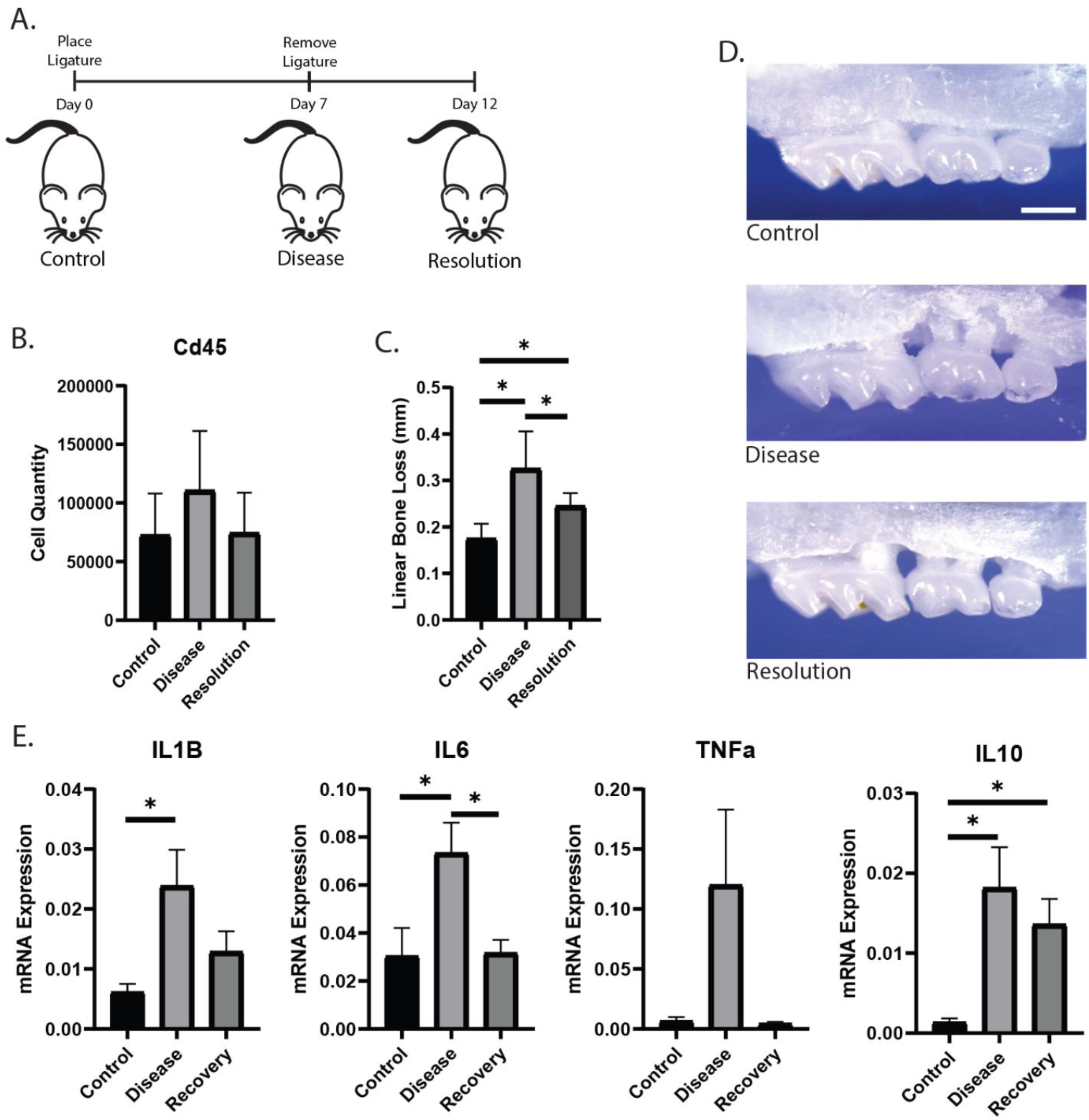
Validation of periodontal disease induction and resolution model. (A) A ligature model of periodontal disease was implemented in adult male mice (C57bl/6, n=7/group), and mice were evaluated in conditions of control, disease, and resolution. (B) Analysis of Cd45+ immune cells isolated from the periodontium via flow cytometry. (C) The maxillary alveolar bone of the mice was evaluated for extent of bone loss across the three conditions. (D) Representative photos of the maxillary alveolar process from mice of each group. Scale bar=1mm. (E) qPCR analysis of gene expression within the maxillary gingival tissue from the maxilla across the three conditions. *p<0.05

### Single-cell RNA sequencing identifies a heterogenous immune cell population isolated from the oral mucosa

To understand the differential cellular immune response across the three conditions of periodontal health, disease, and resolution, we used scRNAseq to analyze Cd45+ immune cells isolated from the periodontium of mice (n=3/group). Cells were isolated via fluorescence-activated cell sorting (FACS), and single-cell library preparations were made using the 10X genomics pipeline. Sequencing was performed on an average of 2,714 cells per mouse with a mean of 202,085 reads per cell. An expanded QC analysis of each sample is presented in Supplemental Figure 1. Figure 2A demonstrates the uniform manifold approximation and projection (UMAP) representation of the immune cells across all conditions. Immune cell types were assigned based on differential expression of known marker genes (Fig. 2B), with each cell type demonstrating a distinct transcriptional profile (Fig. 2C, Supplemental Table 1). We observed a change in the proportion of immune cells isolated from the periodontium across the three conditions (Fig. 2D,E). The disease condition was associated most prominently with an expansion of neutrophils and decrease in the proportion of B cells.

**Figure 2.**
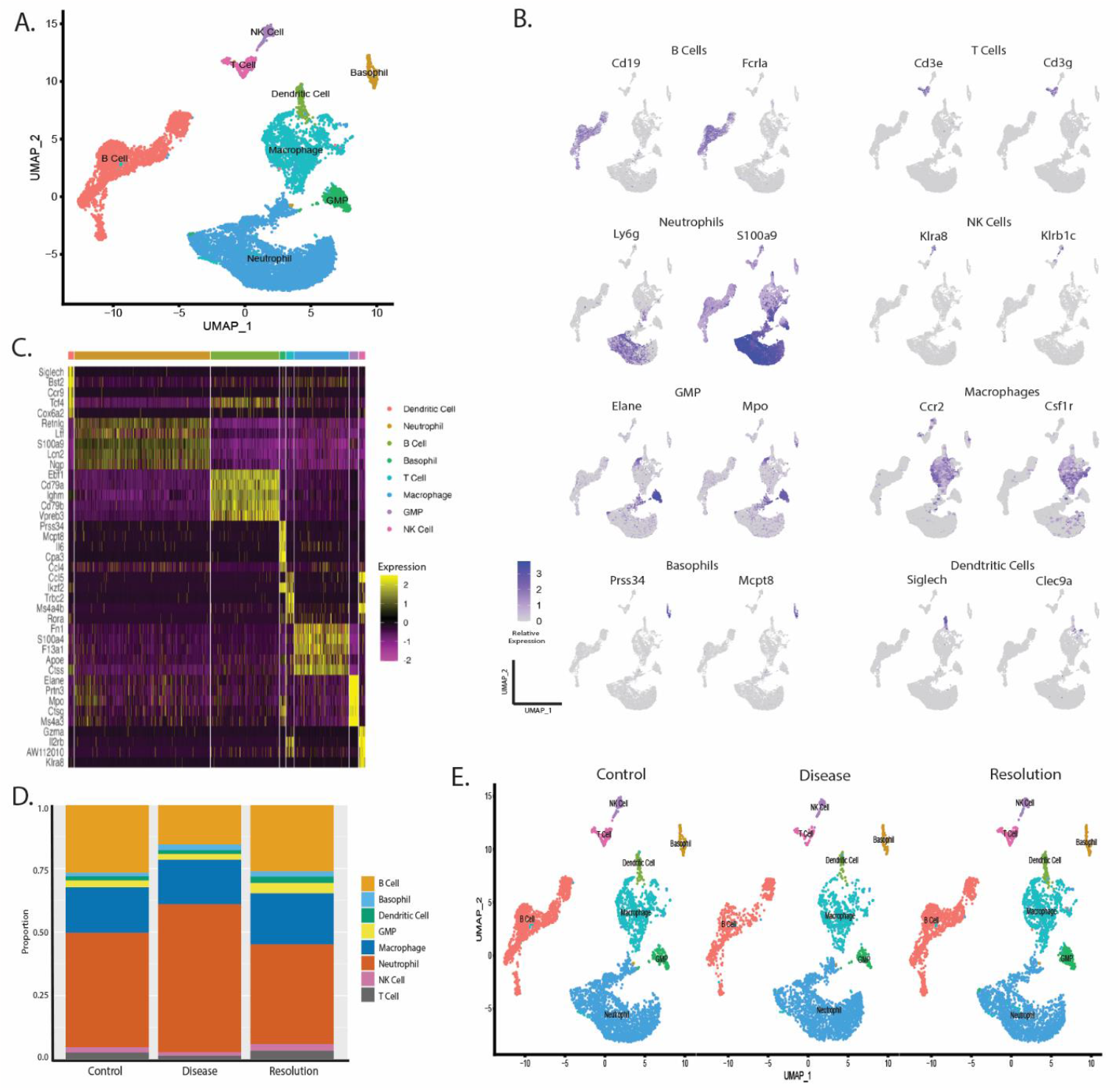
Single-cell RNA sequencing identifies a heterogenous immune cell population isolated from the oral mucosa. scRNAseq analysis was completed on immune cells (Cd45+) isolated from the periodontium of mice in three groups; controls, periodontal disease induced, and periodontal disease resolution (n=3/group). (A) Uniform manifold approximation and projection (UMAP) representation of the immune cells identified in the scRNAseq analysis. (B) Feature plots demonstrate the differential expression of characteristic cell type gene markers used to define immune cell types. (C) Heatmap demonstrates the transcriptional differences across the defined cell types. (D,E) Bar plot and UMAP demonstrates the change of each cell type across the three groups of periodontal control, disease, and resolution. Natural killer cells (NK Cells), Granulocyte-monocyte progenitor cells (GMP).

### Gene signatures shift in immune cells across conditions of periodontal health, disease, and resolution

We next evaluated shifts in gene expression across the three conditions. Differential gene expression analysis was used to compare disease versus control groups and resolution versus disease groups. In this way, we were able to evaluate shifts in gene expression during the change from periodontal health to disease (Disease vs Control) and during the change from periodontal disease to resolution (Resolution vs Disease). Volcano plots demonstrate the significant differentially expressed genes (DEGs) globally across all cell types (Padj<0.05) (Fig. 3A). A total of 237 and 156 genes were significantly up and down regulated respectively in Disease vs Control comparisons, and 192 and 416 genes were significantly up and down regulated respectively in Resolution vs Disease comparisons. Inflammatory genes that have a strong association with periodontal disease pathology, such as Il-1β and Tnfα, were significantly upregulated in Disease vs Control and down regulated in Resolution vs Disease. Similar trends were observed with genes encoding chemokines and matrix metalloproteinases. A full list of the DEGs is presented in Supplemental Table 2. Figure 3B presents the DEGs by cell type. The DEGs are largely enriched within neutrophils and macrophages. In the Resolution vs Disease comparison, DEGs become more apparent in B cells as well. Focusing on the DEGs in neutrophils and macrophages, a strong trend towards the upregulation of genes in Disease vs Control is countered by a downregulation of genes in Resolution vs Disease (Fig. 3B). The heatmap in Figure 3C lists the 148 genes in neutrophils and macrophages that were conversely regulated across the two comparison groups. These 148 genes were a subset of neutrophil and macrophage DEGs presented in Figure 3B that were both upregulated in the Disease vs Control comparison and downregulated in the Resolution vs Disease comparison (Supplemental Table 3). This gene set consists of an enrichment of pro-inflammatory genes upregulated in the pathological inflammatory status of periodontal disease and then downregulated during the resolution of disease. We also observed another subset of DEGs (n=79) that were only upregulated in the Resolution vs Disease comparison (Fig. 3D, Supplemental Table 3). This subset represents, in part, the transcriptional activity uniquely involved with the resolution of inflammation. Gene ontology analysis was further performed on the DEGs significantly up and downregulated in Resolution vs Disease (Fig. 3E). Resolution was associated with an upregulation of genes enriched for apoptotic signaling pathway and pattern recognition signaling. Genes enriched in immune response, cytokine mediated signaling, and response to LPS were downregulated.

**Figure 3.**
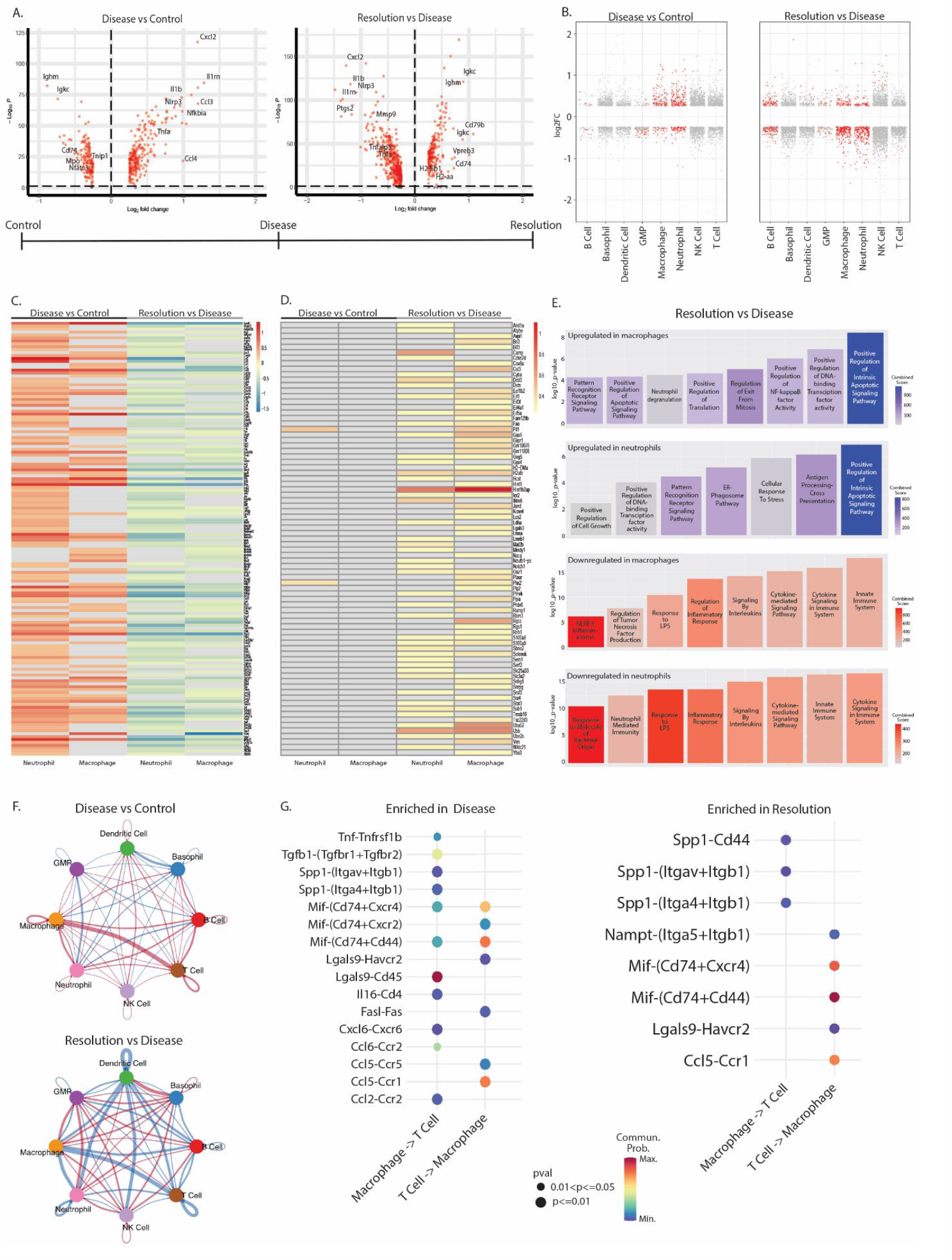
Gene signature shifts identified in immune cells across control, disease, and resolution. (A) Volcano plots demonstrate the total differential gene expression in the diseased group compared to controls (left) and in the resolution group compared to the diseased group (right) (Padj<0.05). (B) Dot plots depict the differential gene expression by cell type in the diseased group compared to controls (left) and in the resolution group compared to the diseased group (right) (Log2FC>0.25, Padj<0.05). (C) Heatmap shows the 148 genes that were inversely regulated across the two comparisons in neutrophils and macrophages. (D) Heatmap shows the 79 genes that were only upregulated in the Resolution vs Disease condition and not differentially regulated in the Disease vs Control condition in neutrophils and macrophages. (E) Gene ontology analysis was performed on the DEGs significantly up and downregulated in Resolution versus Disease comparisons. F) Gene signature shifts were also associated with differential enrichment of cell-cell communication pathways across conditions. Circle plots show the relative quantity of enriched ligand-receptor gene interactions across the immune cell types. The thickness of the line demonstrates the quantity of potential interactions. A red line represents an increase in the number of interaction within the given comparison and the blue line represents a decrease. (G) Dot plots demonstrate the enriched ligand-receptor gene pairs that facilitate macrophage and T cell interactions in conditions of Disease (left) and Resolution (right).

### Gene expression shifts are associated with enriched cell-cell communication pathways across conditions

We also wanted to understand how the shift in gene expression across groups may affect cellular communication between immune cells. We performed cell-cell communication analysis using CellChat to evaluate differential enrichment of ligand-receptor gene pairs. Figure 3F presents circle plots that show the relative quantity of enriched ligand-receptor gene interactions between cell types. The thickness of the line demonstrates the quantity of potential interactions, and the red and blue color represents an increase and decrease respectively of the number of interactions during Disease vs Control (top) and Resolution vs Disease (bottom). An increase in communication was observed between macrophages and T cells during disease compared to control conditions (thick red line). The Resolution vs Disease comparison was associated with an overall decrease in cell-cell communication (blue lines), including decreased communication between macrophages and T cells. The enriched ligand-receptor gene pairs that facilitate macrophage and T cell interactions are shown in Figure 3G. The disease group is characterized by an increased number and diversity of ligand-receptor pairs compared to resolution. Macrophage interaction with T cells was shown to be facilitated through multiple ligand-receptor pairs including the macrophage inhibitory factor (Mif)-Cd74 pathway.

### Immune cell subgroups demonstrate dynamic changes across control, disease, and resolution conditions

We next wanted to better understand the heterogeneity within each cell type and identify potential functional subpopulations of immune cells and their differential expansions across conditions. Unsupervised clustering was performed across all conditions and identified 21 transcriptionally distinct immune cell subgroups (Fig. 4A). Multiple cellular subgroups were identified within the B cell, neutrophil, and macrophage populations. The proportion of the subgroups within each cell type were observed to change across control, disease, and resolution conditions (Fig. 4B-D). We further evaluated the transcriptional profile of each subgroup to gain a better understanding of their activity (Fig. 4B-D, Supplemental Table 4).

**Figure 4.**
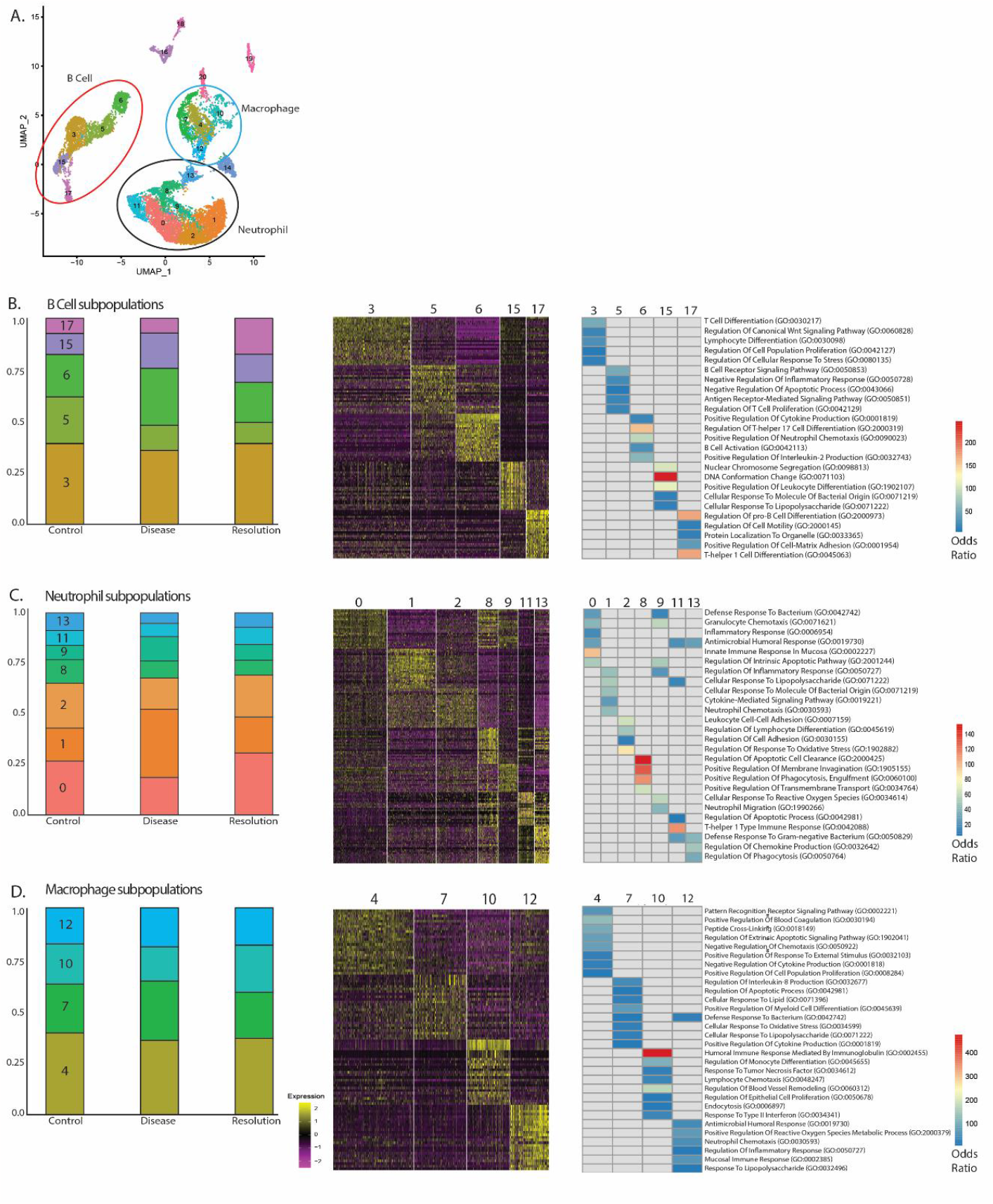
Immune cell subgroups demonstrate dynamic changes across control, disease, and resolution conditions. (A) Unsupervised clustering analysis identified 21 transcriptionally distinct immune cell subgroups across all Cd45+ immune cells analyzed. Multiple subgoups were identified within neutrophils, macrophages, and B cells. Within the (B) B Cells, (C) neutrophils, and (D) macrophage populations, subgroups were analyzed for the proportion of each subgroup across conditions of control, disease, and recovery. Heatmaps and gene ontology analysis demonstrate the differential transcriptional phenotype of each subgroup within the B cell, neutrophil, and macrophage cell types.

In B cells, the resolution condition was associated with an expansion of subgroup 17 (Fig. 4B). This subgroup appeared to be a pro-B cell based on significant differential expression of genes that have been previously shown to be enriched in B cell precursor populations (Vpreb1, Igll1, Lef1)^17,18^. The expansion of this subgroup during the resolution condition suggests B cell recruitment and differentiation within the periodontium is occurring during the resolution of periodontal disease. Also, B cell subgroup 5 was present in higher proportions during the control conditions compared to disease and resolution. Subgroup 5 was enriched for immunoregulatory GO terms such as negative regulation of inflammatory response, negative regulation of apoptotic process, and antigen receptor-mediated signaling pathway. The decreased representation of cells in subgroup 5 during disease and resolution condition represents a decreased immunoregulatory capacity of B cells during those conditions and may position subgroup 5 with a homeostatic role during conditions of periodontal health.

In neutrophils, the proportion of cells in subgroup 1 increased in the disease condition compared to control and resolution conditions (Fig. 4C). The transcriptional profile of subgroup 1 is enriched with expression of pro-inflammatory genes (Il1β, Cxcl2, Ccl3, Csf1, Nfkbia), and GO terms associated with the initiation of inflammation, including cellular response to lipopolysaccharide, cellular response to molecule of bacterial origin, and cytokine mediated signaling pathway. This analysis suggests subgroup 1 contributes to the propagation of inflammation, which is a well described role of neutrophils in periodontal disease. Subgroup 8 maintained a relatively consistent proportion throughout the three conditions. The transcriptional profile of cluster 8 was significant for upregulation of phagocytic and cell clearance GO terms, suggesting a possible homeostatic or regulatory role of this subgroup.

In macrophages, subgroup 7 expanded during the disease condition and demonstrated a proinflammatory transcriptional phenotype, including Mpo, Elane, Ccr2, Prtn3, Ctsg (Fig. 4D). Subgroup 7 was also enriched with inflammatory-associated GO terms including defense response to bacterium, cellular response to oxidative stress, and positive regulation of cytokine production. In contrast, subgroup 10 expanded during the resolution condition. This subgroup had a unique transcriptional profile that was enriched for GO terms that may suggest a critical role of these macrophages in the resolution of periodontal disease. These GO terms include lymphocyte chemotaxis, regulation of blood vessel remodeling, regulation of epithelial cell proliferation, and endocytosis.

### Macrophage subgroup demonstrates resolving phenotype

We wanted to further investigate the resolution-associated phenotype observed in macrophage subgroup 10. Violin plots demonstrate the differential enrichment of genes in subgroup 10 compared to the other macrophage subgroups (Fig. 5A). The genes enriched within this subgroup include Tnip3, Tnfsf9, and Apoe. Tnip3 encodes TNFAIP3 interacting protein 3 which negatively regulates NF-kappa B activation and competes against Tnfα, Tlr3 and Il-1β signaling to downregulate inflammation^19,20^. Tnip3 expression by macrophages has previously been demonstrated as characteristic of an anti-inflammatory macrophage response^21^. Tnfsf9 encodes the TNF superfamily member 9 protein. Tnfsf9 has been shown to induce an anti-inflammatory macrophage phenotype characterized by Il-10 production^22^. Apoe expression by macrophages has been previously characterized in a select immunoregulatory subpopulation^23^, possible acting to dampen T cell activation^24^. In addition, subgroup 10 showed a significant decrease in expression of Fn1 compared to the other subgroups. High Fn1 expressing macrophages have been shown to perturb normal tissue healing and to be involved in pathologic fibrosis in kidney and heart disease and systemic sclerosis^25,26^.

**Figure 5.**
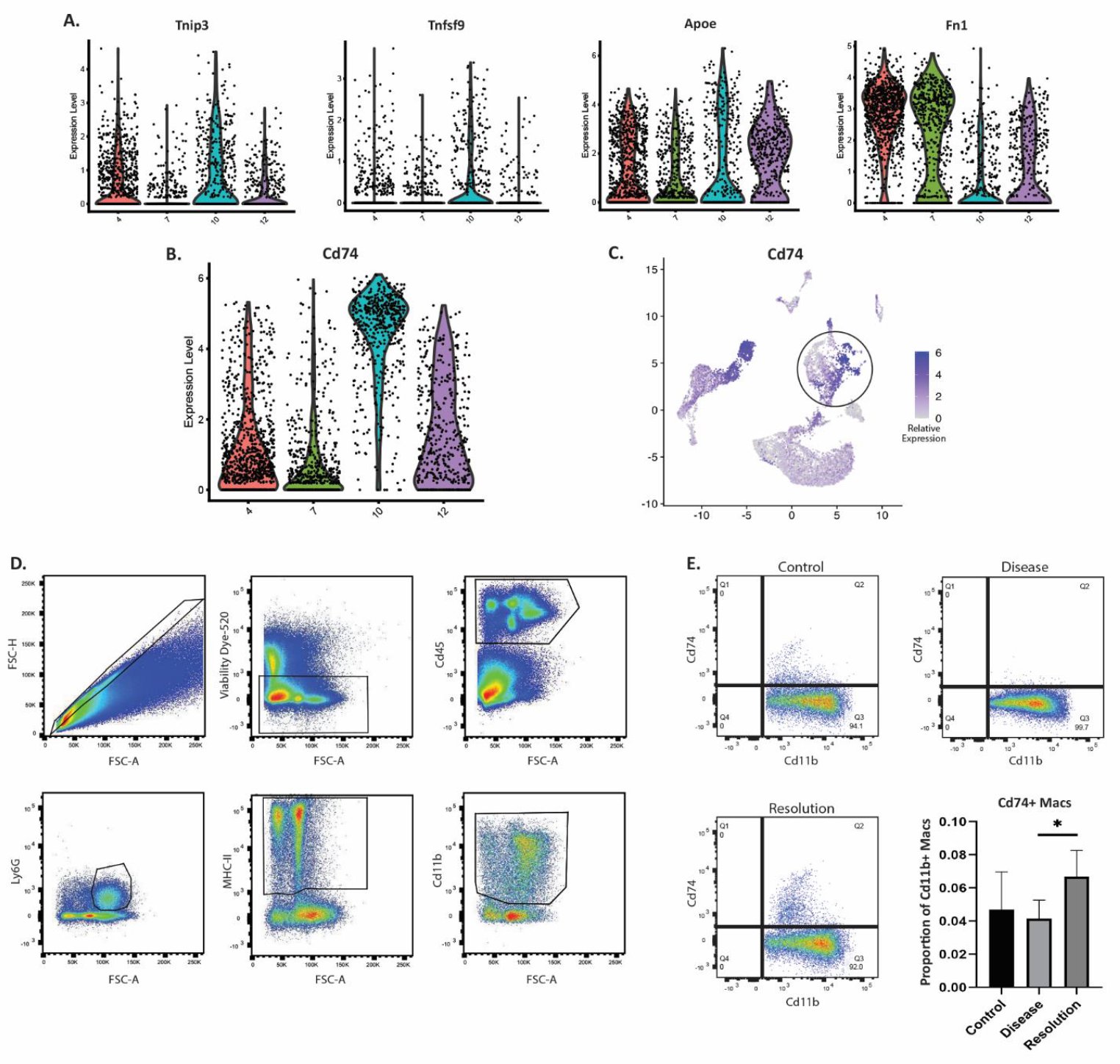
Macrophage subgroup demonstrates resolving phenotype. (A) Violin plots demonstrate the differential expression across all macrophage subgroups of the selected immunoregulatory genes. (B,C) Violin plots and feature plots show that Cd74 expression was highly enriched in subgroup 10 compared to other macrophage subgroups. The presence of a Cd74-expressing macrophage subgroup was validated in vivo. (D) Gating strategy for flow cytometry analysis used to identify Cd74+, Cd11b+ macrophages. (E) Representative quadrant gating applied to samples in each condition (n=7/condition) to identify Cd74+, Cd11b+ macrophages. Bar plot demonstrates the proportion of Cd74+, Cd11b+ macrophages to all Cd11b+ macrophages isolated from the periodontium of mice in control, disease, and resolution conditions.

Subgroup 10 was also transcriptionally defined by the significant upregulation of Cd74 expression compared to the other macrophage subgroups (Fig. 5B). Feature plot shows the differential expression of Cd74 across the macrophage population (Fig. 5C). To further validate the presence of a Cd74+ macrophage population in the periodontium, we used flow cytometry to analyze cells isolated from the periodontium of mice during control, disease, and resolution conditions (Fig. 5D). Here, we validated the presence of Cd11b+, Cd74+ macrophages in the periodontium and showed that the resolution condition was associated with a significant increase in the proportion of Cd74+ macrophages compared to the disease condition (Fig. 5E).

## Discussion

The current study provides an immune cell atlas of the mouse periodontium during three distinct conditions of periodontal health, disease, and resolution. For modeling disease resolution, we induced periodontal disease and then allowed the mice to recover for 5 days after the ligature was removed. Within the 5-day period, the bone loss improved from the disease condition but did not return to base line levels observed in the control condition (Fig. 1). Similarly, inflammatory cytokine expression in the tissue decreased in the resolution group compared to disease, but not all cytokines measured returned to the baseline of the control condition (Fig. 1). Other groups have utilized similar periodontal resolution models in mice and evaluated the extent of healing^27,28^. Using this resolution model, Wong, et al. showed gains in bone levels and decreased osteoclast quantity by 1 week following ligature removal and a near complete return to baseline levels by 2 weeks after ligature removal^28^. Our findings here validate our model of periodontal disease resolution, as the 5-day period post ligature removal was sufficient to observe active resolution of disease without a complete return to baseline control conditions. Therefore, we were confident in our subsequent characterization of the transcriptional processes involved in the resolution of periodontal disease via scRNAseq.

The shift from disease to resolution was associated with a decrease in numerous pro-inflammatory genes across macrophages and neutrophils (Fig. 3). We also observed a subset of genes that was solely upregulated in the resolution condition and represents, in part, the transcriptional processes involved with the active resolution of periodontal disease (Fig. 3D). Gene ontology analysis of this subset showed a significant upregulation of apoptotic signaling pathways in neutrophils and macrophages (Fig. 3E). Apoptosis of immune cells is required for the resolution of inflammation to remove their effects in tissue and prevent further granulocyte release^29,30^.

Our clustering analysis demonstrated multiple cell subgroups in neutrophils, macrophages, and B cells. Interestingly, we did not observe multiple subgroups within T cells. T cell subpopulations, such as Th17 and Tregs, have been well characterized in periodontal health and disease^31^. In our analysis, T cells consisted of 2.5% of the total immune cells isolated from the periodontium. Others have similarly analyzed Cd45+ immune cells isolated from the periodontium of mice via scRNAseq and showed a T cell fraction that was comparable^32^ or markedly greater^33^ to our T cell fraction. Identifying multiple T cell clusters in our data set may have been achieved by performing a second re-clustering analysis using a higher resolution of only T cells. However, others have found that resolving certain T cell subpopulations requires protein characterization and may not be readily achieved with transcriptomic analysis alone^15^.

In addition, we observed multiple transcriptionally distinct macrophage subgroups that differentially expanded or contracted across conditions of periodontal health, disease, and resolution. Others have used scRNAseq to demonstrate pathologic roles of macrophage subpopulations in acute lung inflammation^34^, and have shown that pharmacologically targeting a subpopulation was beneficial in kidney injury^35^. Regenerative or healing macrophage subpopulations have also been identified via scRNAseq to promote osteointegration around titanium tibia implants^36^ and angiogenesis and scar-free healing of skin grafts^37^. To date, there are limited studies that have used scRNAseq to identify macrophage subpopulations in the periodontium and allow the comparisons of these subpopulations across conditions of health, disease, and resolution.

We further focused on macrophage subgroup 10, which was observed to expand during the resolution condition and demonstrated an immunoregulatory transcriptional profile with an enrichment of Tnip3, Tnfsf9, and Apoe and downregulation of Fn1. We also observed Cd74 to be significantly overexpressed in subgroup 10 and we validated Cd74 as a cell marker for this macrophage subgroup via flow cytometry. Flow cytometry analysis showed a distinct population of Cd74+, Cd11b+ macrophages across the three conditions. The Cd74+, Cd11b+ population demonstrated a significant increase during the resolution conditions as was demonstrated via scRNAseq. Cd74 is expressed on multiple cell types including B cells and macrophages^38^, as we have similarly replicated here (Fig. 5C). To only quantify Cd74+ macrophages, we utilized a strict gating strategy that selected against other antigen presenting cells (MHC-ClII) and neutrophils (Ly6G) (Fig. 5D). Cd74 has been shown to have an important role in promoting healing and limiting inflammatory damage of heart, kidney and lung tissue following injury or disease^39^. However, the role of Cd74 in the resolution of periodontal disease has not been previously described. Cd74 has been most prominently studied as the receptor for the macrophage migration inhibitory factor (MIF)^40^. Interestingly, our cell-cell communication analysis showed a Cd74-MIF interaction mediating macrophage-T cell interactions (Fig. 3G).

This study did not fully define the functional role of a Cd74+ macrophage subpopulation in the resolution of periodontal disease. However, we were successful in validating Cd74 as a marker for a macrophage subgroup with an immunoregulatory transcriptional profile that expanded during resolution. These findings are important for future investigations that further characterize this subgroup and identify potential therapeutics that target its expansion in the promotion of inflammatory resolution. Further, the large cell atlas generated here should provide a resourceful dataset to compare immunological changes across periodontal health, disease, and resolution.

## Author Contributions

Cantalupo P, contributed to data acquisition, analysis, and interpretation, drafted and critically revised the manuscript; Diacou A, Park S, Chen J, Glenn D contributed to data acquisition and analysis; Soman V and Chandran U contributed to data analysis and interpretation and critically revised the manuscript. Clark D contributed to conception, design, data acquisition, analysis, and interpretation, drafted and critically revised the manuscript.

## Data availability Statement

The single cell RNA sequencing data reported in this article has been uploaded to NCBI GEO and the accession number can be provided upon request.

---

**Supplemental Figure 1.**
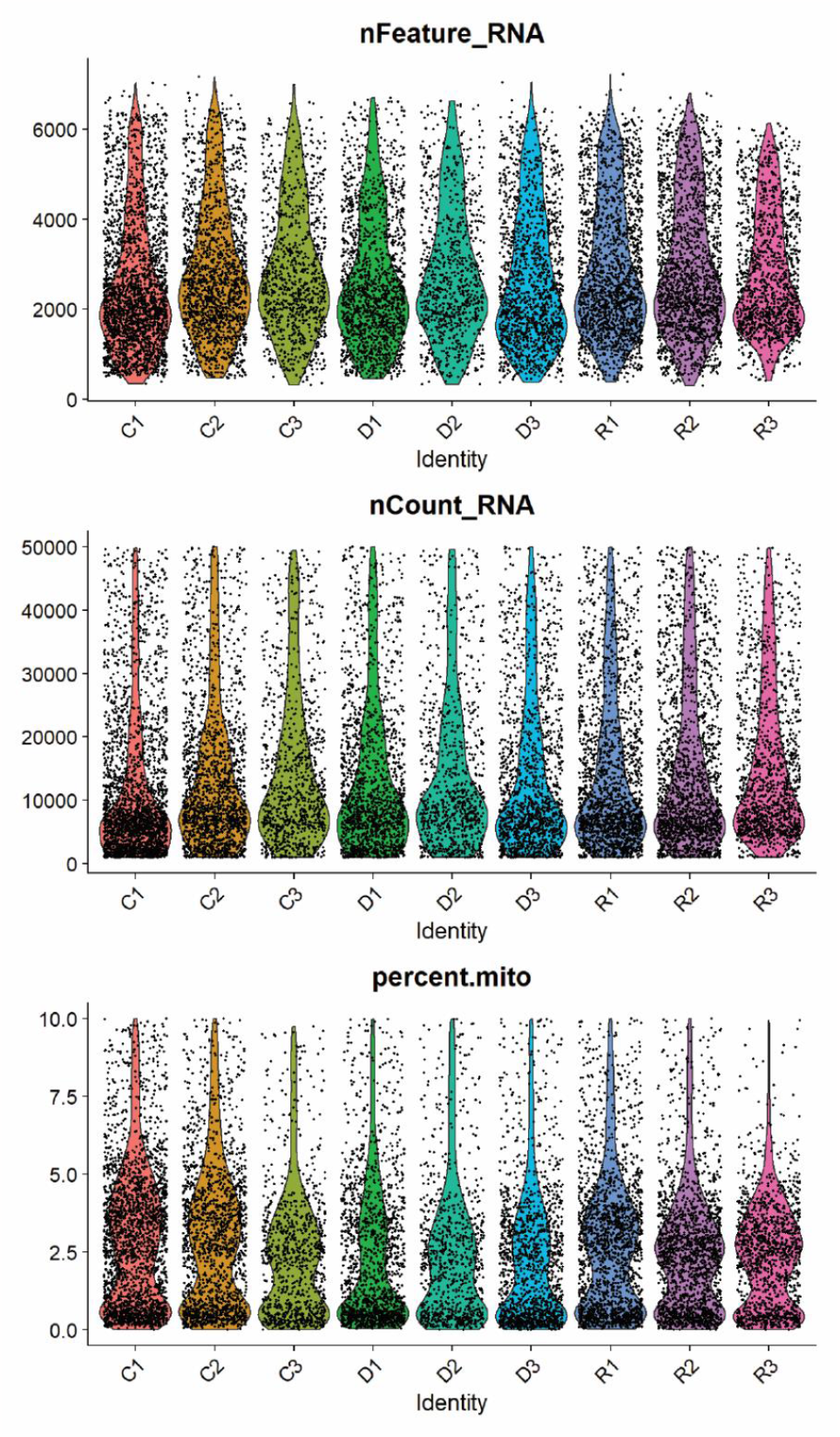
Quality control analysis of each sample from control (C), disease (D), and resolution (R) conditions. Violin plots shows the number of detected genes for each cell (nFeature_RNA), the number of detected unique molecular identifiers (UMI) for each cell (nCount_RNA), and the percentage of mitochondrial genes for each cell (percent.mito).

